# Topoisomerase 2β induces DNA breaks to regulate human papillomavirus replication

**DOI:** 10.1101/2021.01.07.425831

**Authors:** Paul Kaminski, Shiyuan Hong, Takeyuki Kono, Paul Hoover, Laimonis Laimins

## Abstract

Topoisomerases regulate higher order chromatin structures through the transient breaking and re-ligating of one or both strands of the phosphodiester backbone of duplex DNA. TOP2β is a type II topoisomerase that induces double strand DNA breaks at topological-associated domains (TADS) to relieve torsional stress arising during transcription or replication. TADS are anchored by CTCF and SMC1 cohesin proteins in complexes with TOP2β. Upon DNA cleavage a covalent intermediate DNA-TOP2β (TOP2βcc) is transiently generated to allow for strand passage. The tyrosyl-DNA phosphodiesterase TDP2 can resolve TOP2βcc but failure to do so quickly can lead to long-lasting DNA breaks. Given the role of CTCF/SMC1 proteins in the HPV life cycle we investigated if TOP2β proteins contribute to HPV pathogenesis. Our studies demonstrated that levels of both TOP2β and TDP2 were substantially increased in cells with high risk HPV genomes and this correlated with high amounts of DNA breaks. Knockdown of TOP2β with shRNAs reduced DNA breaks by over 50% as determined through COMET assays. Furthermore this correlated with substantially reduced formation of repair foci such as γH2AX, pCHK1 and pSMC1 indicative of impaired activation of DNA damage repair pathways. Importantly, knockdown of TOP2β also blocked HPV genome replication. Our previous studies demonstrated that CTCF /SMC1 factors associate with HPV genomes at sites in the late regions of HPV31 and these correspond to regions that also bind TOP2β. This study identifies TOP2β as responsible for enhanced levels of DNA breaks in HPV positive cells and as a regulator of viral replication.

## Introduction

Human papillomaviruses (HPVs) infect stratified epithelia and link their productive life cycles to epithelial differentiation (1, 2). HPVs infect cells in the basal layer that become exposed through microwounds and establish their double stranded circular DNA genomes in the nucleus as low copy episomes at 50 to 100 copies per cell. Productive viral replication or amplification only occurs in highly differentiated suprabasal cells and is dependent upon activation of the ataxiatelangiectasia mutated kinase (ATM) (3) as well as the ataxia-telangiectasia and RAD3-related kinase (ATR) pathways (4–6). These pathways are activated by the action of viral proteins, such as E6 and E7, alone and do not require viral replication (3–5). Previous studies have shown that activation of these pathways in HPV positive cells occurs through induction of high levels of DNA breaks in both viral and cellular DNAs (7). The breaks in viral genomes are rapidly repaired by DNA damage repair pathways and this is necessary for differentiation-dependent genome amplification (7). The mechanism by which HPV proteins induce high levels of DNA breaks, however, has not been clarified.

DNA breaks can be induced by exposure to exogenous DNA damaging agents or through endogenous pathways such as through the action of topoisomerases. Topoisomerases regulate higher order chromatin structures through the transient breaking and re-ligation of one or both strands of the phosphodiester backbone of duplex DNA (8, 9). Type I topoisomerases (TOP1) cleave a single strand of duplex DNA, while type II topoisomerases (TOP2α and TOP2β) generate DNA double strand breaks (DSBs) (9). TOP2α is expressed primarily in proliferating cells while TOP2β is ubiquitously expressed including during differentiation (8, 10). During breakage and re-ligation by TOP2β a covalent intermediate DNA-protein crosslink (TOP2βcc) is generated, which allows for strand passage and/or DNA unwinding to occur while the strand break remains bookmarked by the topoisomerase (11, 12). A certain proportion of TOP2βcc, however, result in long lasting DNA breaks. These complexes can be resolved through the action of TDP2 enzymes (13) or the Mre11/RAD50/Nbs1 (MRN) complexes from the ATM pathway (14).

TOP2β cleavage of DNA is required for proper replication and transcription. Chromatin is arranged in DNA loops called topological-associated domains (TADS) which are anchored by CTCF factors in association with SMC1 cohesin proteins (15). TOP2β associates with CTCF and SMC1 at these anchor sites and this is required to dissipate torsional stress arising from transcription or replication through the formation of DNA breaks (12, 16). Previous studies demonstrated that CTCF/SMC1 binding sequences are present in the late regions of almost all HPVs (17). Importantly knockdown of either CTCF or SMC1 in HPV31 positive cells reduced viral transcription and blocked viral amplification (17). We therefore investigated if TOP2β proteins played a role in regulating HPV pathogenesis and found that levels were increased in cells with high risk HPV genomes and this correlated with increased levels of DNA breaks. Importantly, knockdown of TOP2β reduced the amount of DNA breaks by over 50% which correlated with impaired activation of DNA damage repair pathways. Furthermore, HPV replication was also blocked following knockdown of TOP2β. Our studies identify a major source of DNA breaks in HPV positive cells that is responsible for activation of DNA damage repair pathways and for regulation of viral replication.

## Results

### Levels of TOP2β are increased in HPV positive cells

To investigate whether TOP2β played a significant role in the HPV life cycle, we first examined the levels in cells that maintain either HPV16 or HPV31 episomes by western analysis. The levels of TOP2β were found to be increased by approximately 3 to 5-fold in cells with HPV 16 or 31 episomes (3) in comparison to normal human keratinocytes (Figure 1A). HFK-16 and HFK-31 are stable cell lines that maintain viral episomes and were generated by transfection of human foreskin keratinocytes (HFKs) with recircularized viral genomes (18). CIN 612 cells maintain HPV 31 episomes and were derived from a biopsy of a CIN II lesion (19). To investigate if the levels of TOP2β changed upon differentiation HPV positive cells were induced to differentiate by addition of high calcium media and examined by western analysis. Similar levels of TOP2β were observed in undifferentiated and differentiated HPV positive cells. While HFKs exhibited low levels in undifferentiated cells as well as following 48 hours of calcium-induced differentiation, an increase was consistently observed at 72 hours of differentiation which is after amplification and late gene expression have been completed. (Figure 1B). The reason for this increase is unclear. We also investigated if the increase of TOP2β in HPV positive cells occurred at the level of transcription or post transcriptionally. An analysis of TOP2β transcripts from undifferentiated HFKs and CIN 612 cells showed similar levels indicating regulation occurred at the post-transcriptional level (Figure 1C).

**Figure 1:**
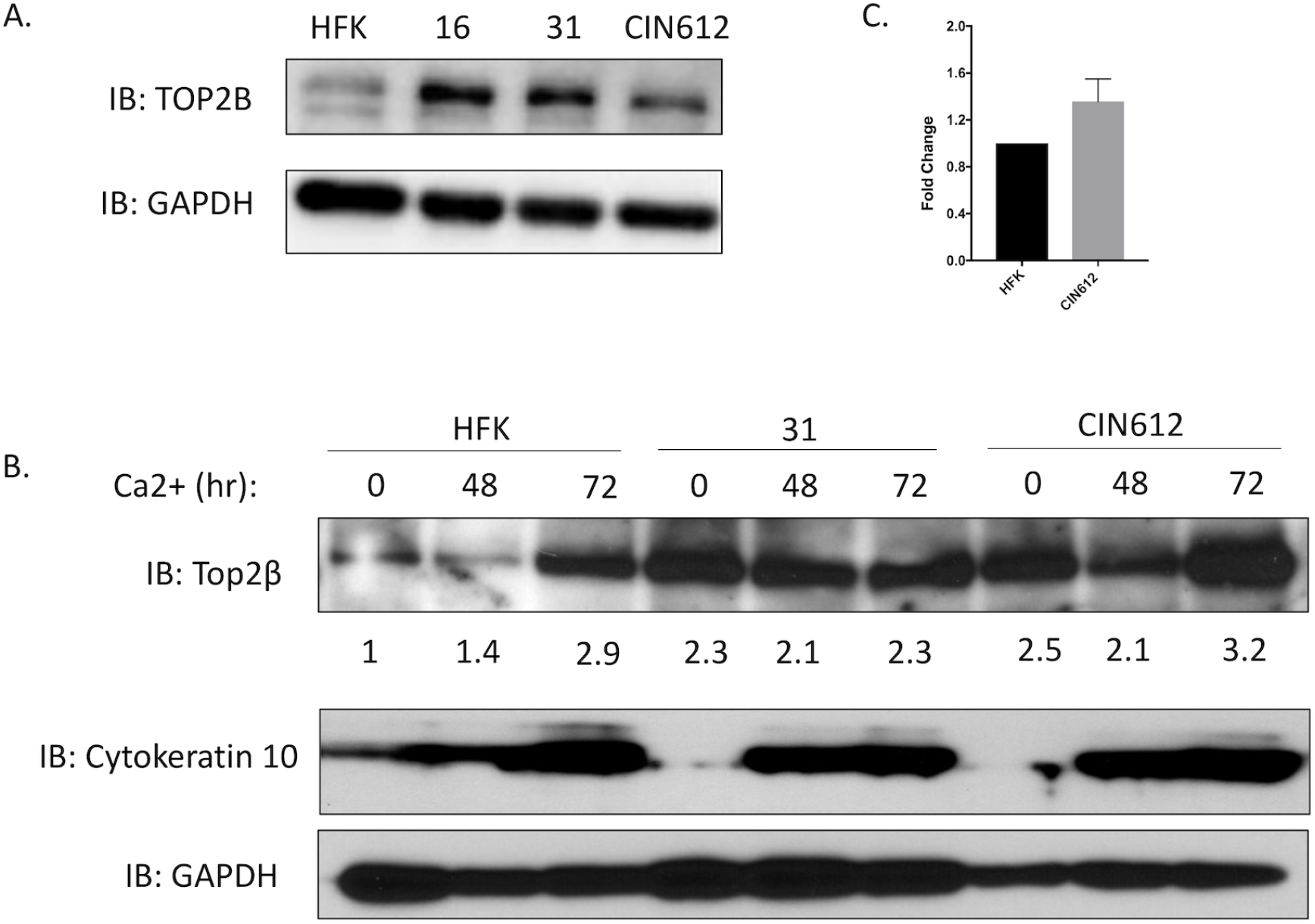
(A). TOP2β levels are increased in HPV positive cells. Monolayer cultures of normal human foreskin keratinocytes (HFKs) along with cells with HPV episomes: HFK-16 (HPV 16 positive); HFK-31 (HPV31 positive) and CIN 612 (HPV 31 positive) were examined by western for levels of TOP2β. GAPDH controls are shown below. Similar results were seen in three separate experiments. (B). TOP2β levels in HPV31 positive cells remain high upon differentiation in high calcium media. HFK, HFK-31 and CIN 612 cells are shown at 48 and 72 hours after calcium switch. Quantitation of band intensities is shown below. Levels of differentiation marker cytokeratin 10 are shown along with GAPDH controls. (C). q-RT-PCR analysis of TOP2β transcripts in undifferentiated HFKs and CIN612 cells

We next investigated if replication of viral episomes or expression of viral proteins alone was responsible for inducing high levels of TOP2β. For this analysis we infected HFKs with retroviruses expressing HPV16 or 31 E6, E7 or the combination of E6 and E7 and isolated stable cells. Examination of extracts from these cells by western analysis demonstrated that E7 was largely responsible for increased levels of TOP2β. (Figure 2). The levels of TOP2β in E7 expressing were similar to those seen in cells that stably maintain viral episomes. We conclude that expression of E7 alone is sufficient to induce high levels of Top2β and viral replication is not required.

**Figure 2:**
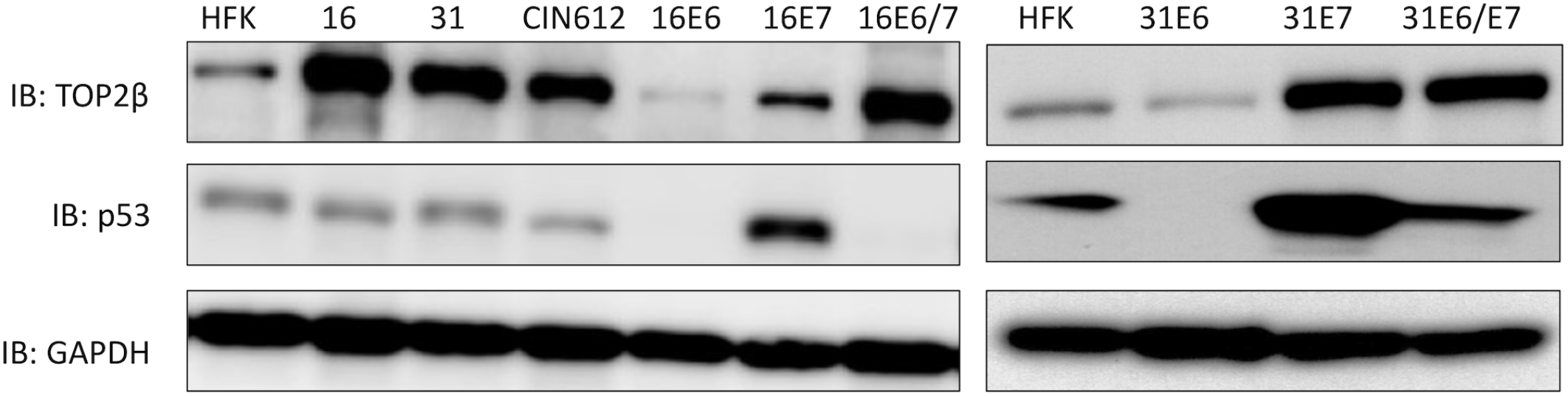
HPV E7 protein is responsible for increased levels of TOP2β. Western analysis of Top2β and p53 in the following cells: HFKs; HFK-16; HFK-31; CIN 612; HPV16 E6; HPV16 E7; HPV16 E6/E7; HPV31 E6; HPV31 E7; HPV31 E6/E7. Cell lines expressing individual E6 or E7 proteins were generated by stable infection of HFKs with retroviral expression vectors expressing the various proteins. GAPDH loading control was included.

### Knockdown of TOP2β impairs HPV replication

To determine if increased levels of TOP2β had an effect on viral functions, we examined the consequences of shRNA knockdowns on HPV replication. CIN 612 cells were first transiently infected with five individual lentiviruses expressing different shRNAs against TOP2β and we identified two that had an effect. While expression of shRNA #3 resulted in a moderate suppression of TOP2β levels, shRNA#4 substantially reduced levels (Figure 3). In addition, knockdown of TOP2β had no significant effect on differentiation, as indicated by expression of cytokeratin-10. Next CIN 612 cells were infected with these two shRNAs, stable lines isolated and screened for the presence of viral episomes in both in undifferentiated cells and following differentiation in high calcium media for 72 hours. As shown in Figure 4, knockdown of TOP2β by either shRNA reduced the levels of viral episomes in undifferentiated cells. Screening for episomal genomes using an exonuclease V resistance assay that identifies circular genomes demonstrated a reduction of 3 to 4 fold in the levels of episomes in undifferentiated TOP2β knockdown cells (Figure 3B). Since episomal templates are required for differentiation-dependent amplification this process was also inhibited. In addition, knockdown of TOP2β had no effect on proliferation as cell lines were passaged over 5 times in culture and grew similar to parental CIN 612 cells. Furthermore early transcripts encoding E7 were detected using q-RT-PCR in both wildtype, control and knockdown cells though levels were reduced by approximately 50% in the latter (Figure 3C). These experiments indicate that TOP2β is a critical regulator of viral replication and may act in part through effects on HPV transcription.

**Figure 3:**
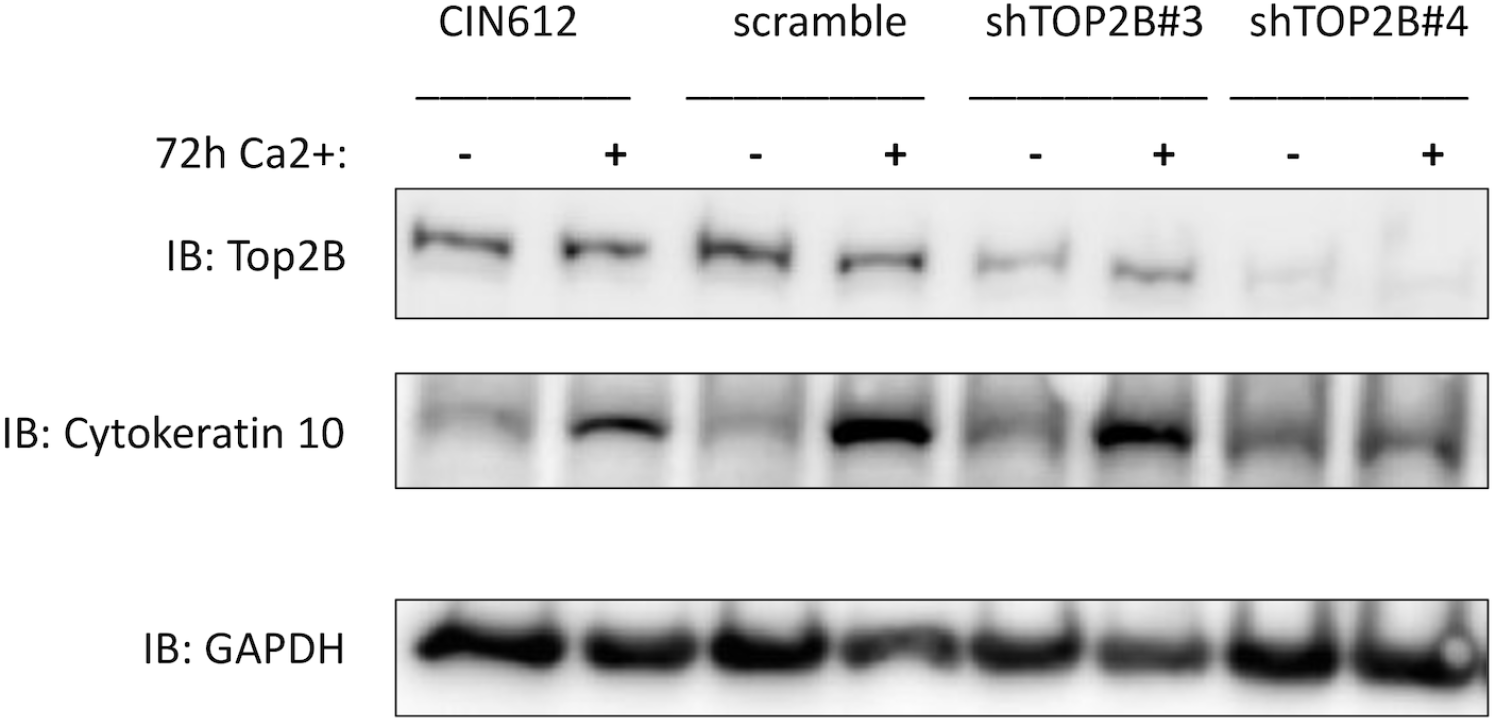
Identification of shRNA constructs that knockdown or reduce TOP2β levels. Lentiviruses expressing either of two shRNAs against TOP2β or a scramble control were generated in 293TT cells and resulting lentiviruses used to infect CIN 612 cells. Following selection and expansion, stable cell lines were isolated. Following differentiation in high calcium media for 0 or 72 hours, cell extracts were examined for levels of TOP2β or cytokeratin 10 by western analysis.

**Figure 4:**
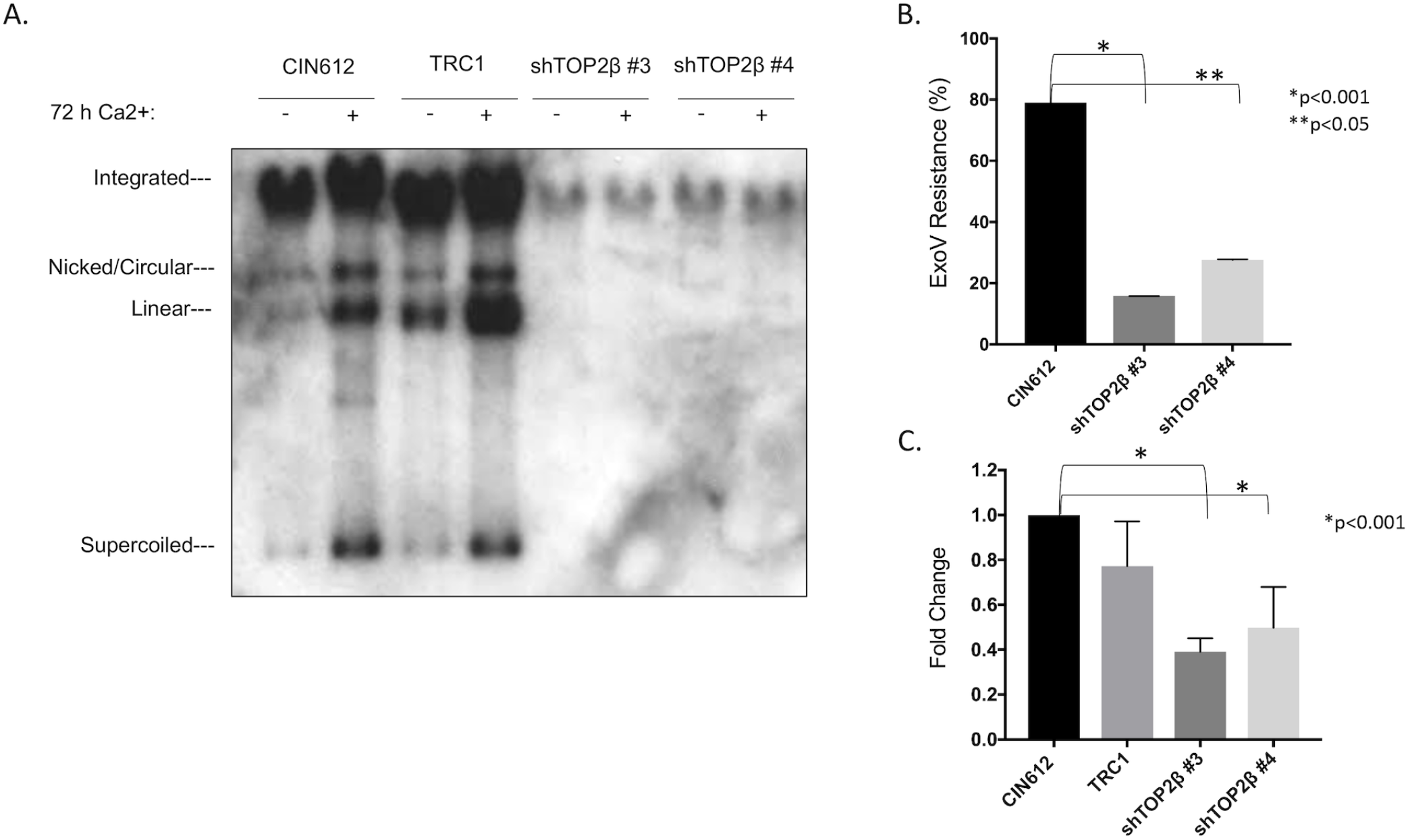
Knockdown of TOP2β in CIN 612 impairs stable maintenance of HPV 31 episomes and amplification upon differentiation. (A). Southern analysis of CIN 612 wildtype and scramble control compared to cells with stable knockdown of TOP2β. Undifferentiated and after 72 of differentiation. Supercoiled episomes, nicked circular, linear, multimers and integrated genomes are indicated. (B). Exo V analysis of episomal forms in undifferentiated CIN 612 cells and stable TOP2β knockdowns P values *p<0,001, **p<0,05 (C). q-RT-PC analysis of HPV early transcripts encoding E7 in undifferentiated cells. Parental CIN 612 cells, shRNA control TRC1 infected CIN 612 cells, stable knockdowns of TOP2β in CIN612 background with shRNA #3 and shRNA #4. P values *p< 0.001

### TOP2β binds to HPV genomes at multiple sites

To begin to understand how TOP2β regulated HPV replication we investigated if it associated viral genomes by performing chromatin immunoprecipitation analyses. For this study, several regions of the HPV31 genome were examined along with binding to cellular ALU sequences that were chosen as representative of cellular sequences. ALU were chosen as they are highly transcribed and present in multiple copies. TOP2β was found to bind at high levels to the L2, E2 ORF and E7 ORF regions of the HPV genomes comparable to that detected at ALU sequences. In contrast, only a low level of TOP2β binding was detected at the URR of HPV 31 (Figure 5). Interestingly, the regions of the HPV31 genome that were found to bind TOP2β are those that contain putative or documented CTCF/SMC1 binding sites (17). No CTCF sites were identified in the URR that also fails to exhibit TOP2β binding. This suggests TOP2β associates to regions of the viral genomes that may be sites for CTCF/SMC1 mediated DNA looping.

**Figure 5:**
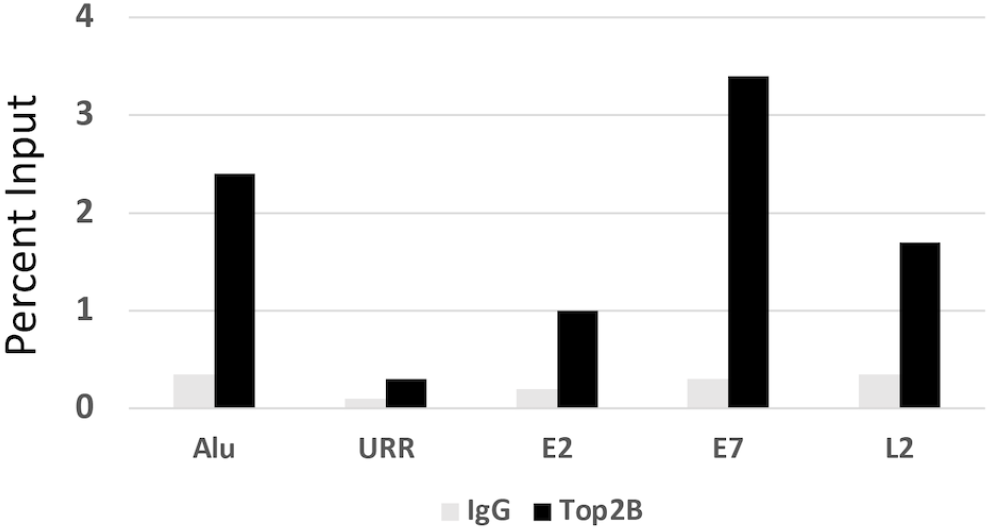
Chromatin immunoprecipitation analysis of TOP2β binding to regions on HPV 31 episomes in comparison to Alu sequences. ChIP analyses were performed on monolayer cultures of CIN 612 cells. Similar results were observed in three independent experiments.

### High levels of TDP2 are present in HPV positive cells

Previous studies have shown that the levels of DNA breaks in HPV positive cells are 2 to 5 fold higher than in normal keratinocytes (7). TOP2β can induce DNA breaks through the formation of TOP2βcc intermediates that are resolved through the action of the tyrosyl-DNA phosphodiesterase TDP2. TDP2 quickly resolves TOP2βcc allowing for re-ligation of the DNA break. We investigated if high levels of TOP2β correlated with increased amounts of TDP2. As shown in Figure 6, TDP2 levels are increased by 5 to 7 fold in cell lines that maintain episomes of HPV 16 or 31 indicating that breaks induced by TOP2β can be rapidly resolved. We next investigated if the increased amounts of TOP2β in HPV positive cells also correlated with high levels of DNA breaks. To distinguish TOP2β induced breaks from those caused by other mechanisms we examined CIN 612 cell lines in which TOP2β levels were decreased by shRNAs with COMET assays. In COMET assays single cells are suspended in agarose on glass slides followed by lysis, electrophoresis and imaging by fluorescence (7). Large unbroken DNAs are retained in nucleoid body while broken DNAs migrate into the tail region. The levels of DNA breaks can be quantified by measuring the signal in the tail relative to that in the head. Examination of wildtype CIN 612 cells or those stably infected with lentiviruses expressing a scrambled control shRNA demonstrated the presence of high levels of DNA breaks (Figure 6B). In contrast, knockdown of TOP2β resulted in a statistically significant decrease in the amount of DNA breaks by approximately 50 to 60%.

**Figure 6:**
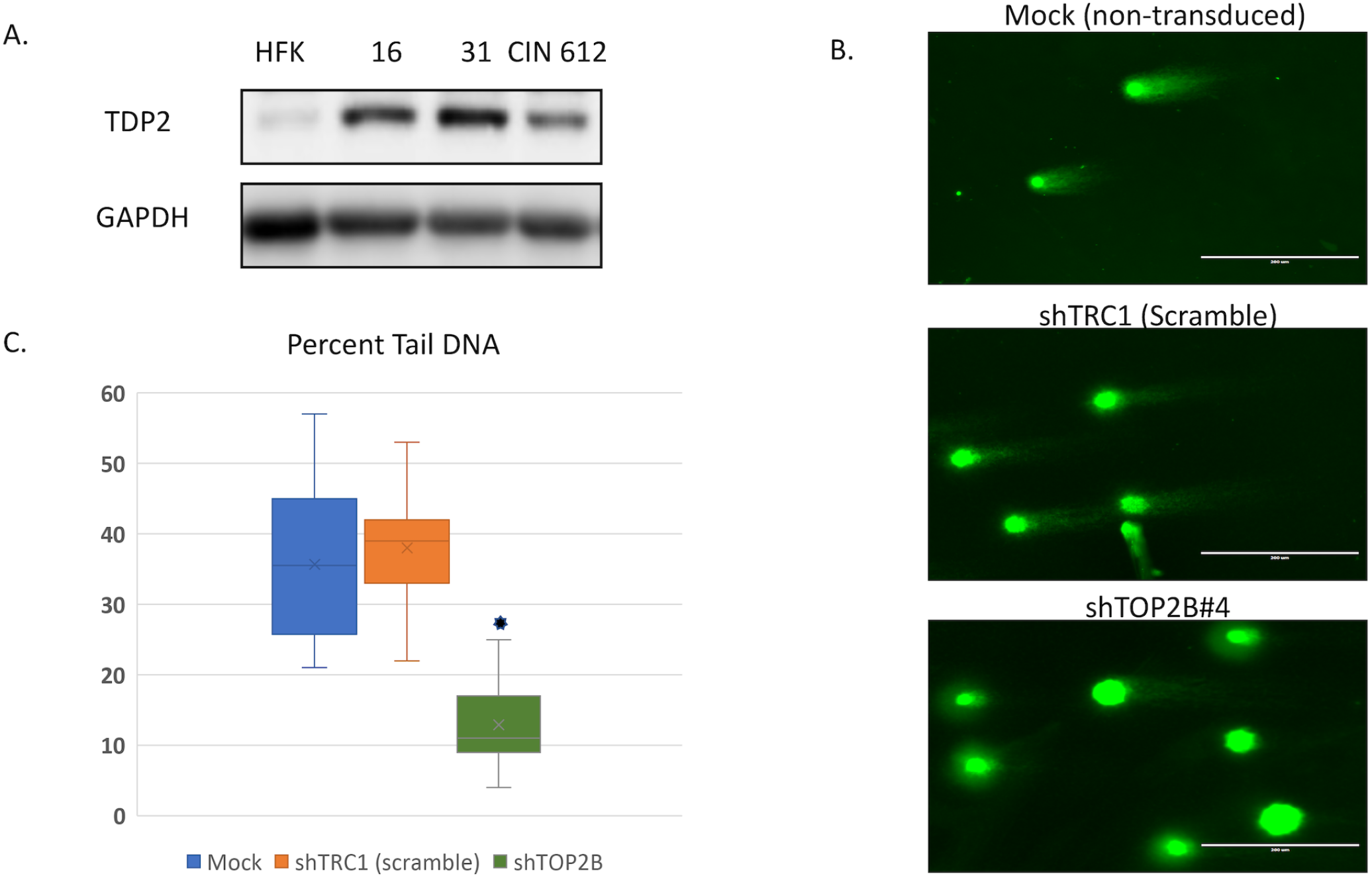
A). TDP2 levels are increased in HPV positive cells. Monolayer cultures of normal human foreskin keratinocytes (HFKs) along with cells with HPV episomes: HFK-16 (HPV 16 positive); HFK-31 (HPV31 positive) and CIN 612 (HPV 31 positive) were examined by western for levels of TDP2. GAPDH controls are shown below. (B). Neutral COMET assays for DNA break formation for in CIN 612 cells with mock, shTRC1 (scramble) and shRNA against TOP2β. Quantitation of percent tail versus nucleoid body shown in graph. A reduction in levels of DNA breaks of greater than 50% was detected in shRNA knockdown cells and is statistically significant. P value *p<0.001

### Knockdown of TOP2β reduces nuclear foci containing DNA damage repair factors

Since the number of DNA breaks was reduced in HPV positive cells upon TOP2β knockdown, we investigated whether this impacted the activation of DDR pathways. HPV activates both ATM and ATR factors and recruits these proteins to nuclear repair foci. To investigate if DDR activation was altered by TOP2β knockdown we examined if there was a reduction in the number of cells with greater than 3 foci comparing CIN 612 wildtype to TOP2β knockdown cells. Immunofluoresence assays indicate that the number of cells with pCHK1, pSMC1 and γH2Ax positive foci was reduced by 50 to 80% in knockdown cells (Figure 7). Similarly the levels of nuclear focus formation of other DDR factors, such as pCHK2, FANCD2, and RAD51, were also reduced. TOP2β nuclear staining was present in a diffuse punctate pattern and reduced in knockdown cells. We conclude that TOP2β activity impacts the recruitment of DNA damage repair factors to foci consistent with the reduced numbers of breaks in the cells.

**Figure 7:**
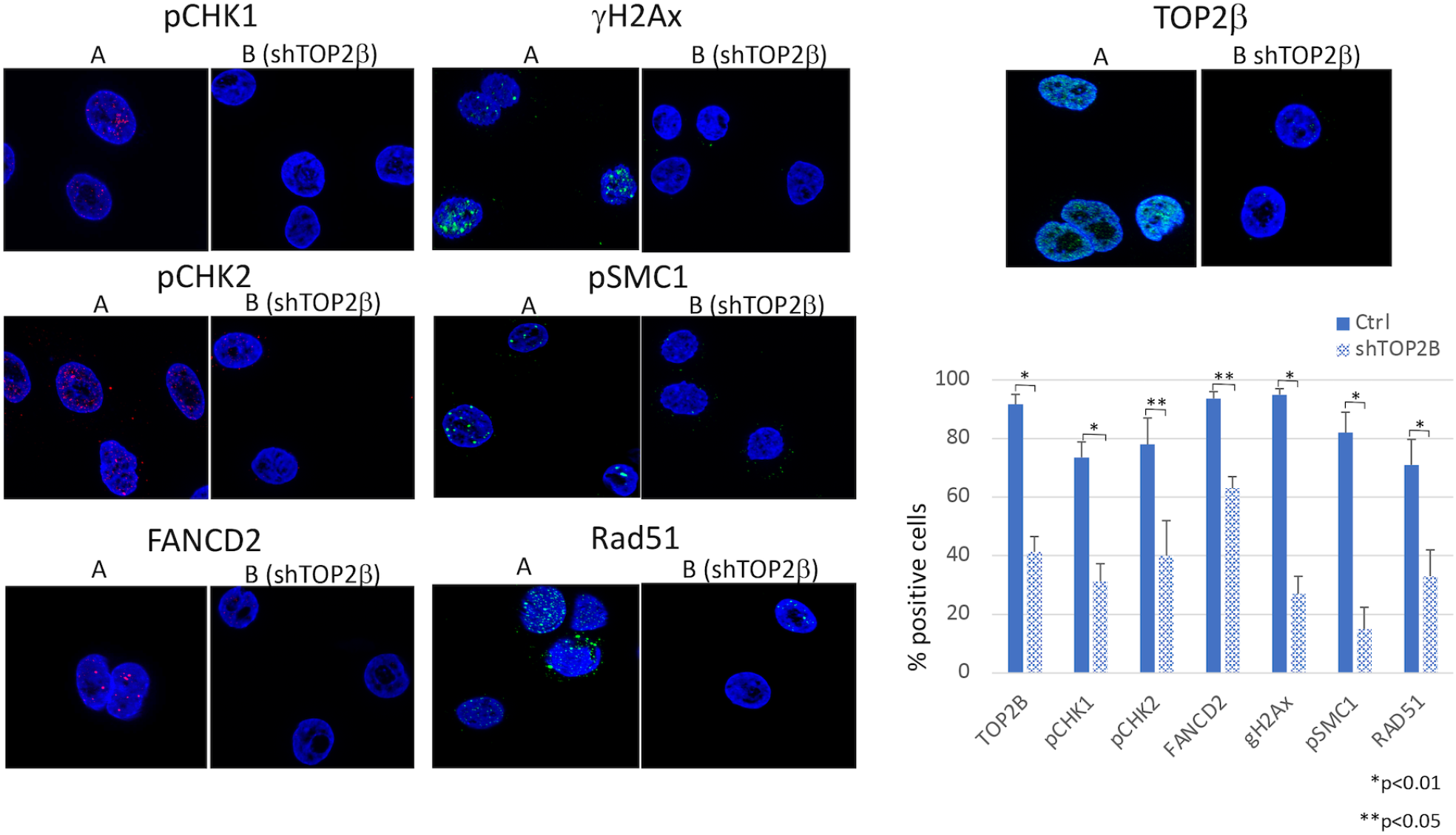
(A) Representative Immunofluorescence staining of TOP2β and other DNA damage factors in CIN612 control and TOP2β knock down cells. High magnification (x63) is shown and indicates the presence of positive foci. (B) Comparative analysis of the percentages of cells that exhibited more than 3 focal localization of DDR factors between CIN 612 control cells and TOP2β knock down cells. P values *p <0.01, **p<0.05

## Discussion

TOP2β relieves torsional stress caused by transcription and replication by inducing DNA breaks followed by re-ligation (11, 16). Our studies indicated that TOP2β levels were elevated in HPV positive cells by three to five fold which can lead to TOP2βcc induced DNA breaks. Our studies further suggest that the increased expression of TOP2β was responsible for over half of the DNA breaks in HPV positive cells and was important for the activation of DNA damage repair pathways. This is particularly significant in HPV positive cells where activation of DNA damage repair pathways is necessary for HPV replication and differentiation-dependent amplification (7). Importantly TOP2β was shown to be important for the stable maintenance of episomes as well as differentiation-dependent amplification which requires the presence of episomal genomes.

Topological-associated domains (TADS) are anchored through the binding of CTCF factors in association with SMC1 cohesin proteins and TOP2β. Previous studies demonstrated that CTCF/SMC1 factors associate with HPV genomes at conserved CTCF binding sites present in the late regions of all HPVs (17). Knockdown of either of these factors with shRNAs blocks HPV31 genome amplification and reduces late gene expression. The HPV 31 L2, L1 and E2 ORFs contain consensus CTCF sequences and our recent 4C chromatin capture analyses demonstrated the formation of DNA loops between these sites as well as a putative cryptic site in the E7 ORF (Mehta, K., Hoover, P. and Laimins, L. unpublished). No CTCF directed looping was detected to sequences in the URR. In the present work our TOP2β chromatin immunoprecipitation analyses showed TOP2β binding to these same 4 regions with no binding was detected at the URR. This is consistent with TOP2β forming complexes with CTCF/SMC1 proteins at CTCF sites in HPV genomes. The Parish laboratory has shown that the CTCF site in HPV18 E2 is critical for viral expression (21) and we have shown that knockdown of CTCF interferes with viral expression (17). These findings indicate that while TOP2β is responsible for global increases in DNA breaks in HPV positive cells, it may have additional activities in directly regulating viral transcription and replication resulting from its binding to HPV genomes. The observation that expression of early viral genes was reduced by 50% upon stable knockdown of TOP2β is consistent with this hypothesis, however, many factors can influence viral expression in cells following integration of HPV genomes into host chromosomes. A more detailed analysis examining expression of all viral transcripts after transient knock down of TOP2β is required for a more complete understanding. In addition, TOP2β or other topoisomerases may play a critical role in helping resolve concatemeric viral genomes resulting from amplification. Amplification has been reported to occur through several potential mechanisms including a rolling circle type process or a novel unidirectional replication scheme involving multiple initiation sites (22, 23). It also is possible that amplification occurs though bi-directional theta structures. All three models require the action of topoisomerases such as TOP2β, TOP2α or TOP1 to resolve concatamers. Interestingly the Androphy lab has reported that E2 associated with TOP1 to facilitate its recruitment to viral episomes for replication and may also facilitate resolution of concatemers (24). TOP2β along with other topoisomerases may thus have multiple functions in regulating the HPV life cycle

Along with increased amounts of TOP2β, high levels of the phosphodiesterase TDP2 (13) which rapidly resolves TOP2βcc structures were also detected in HPV positive cells. This suggests that once generated TOP2βcc are rapidly resolved by TDP2 which is consistent with our previous observations that while DNA breaks are generated in viral genomes they are rapidly repaired (7). Interestingly TOP2βcc has been suggested to play a positive role in regulating pathogenesis of some viruses. Treatment of neurons with etoposide, which stabilizes TOP2βcc, results in increased expression of over 13 genes particularly in the neuronal early response pathway as well as decreased expression of 300 other genes (25). Furthermore, high levels of TOP2βcc, are induced in HSV infected neurons and these are needed for the maintenance of viral latency (26). Whether TOP2βcc structures contribute in a similar manner to the regulation of late HPV functions is an area for future study.

Our studies indicate that TOP2β is responsible for inducing many but not all DNA breaks in HPV positive cells. The question arises as to what other mechanisms could be responsible for causing the other DNA breaks. One possibility is that TOP2α, which is highly homologous to TOP2β, could play a role. TOP2α is expressed primarily in cycling cells where it functions to relieve torsional stress resulting from replication and transcription (10). TOP2α also forms TOP2αcc covalent intermediates that result in DNA break formation (10). In contrast to TOP2β. TOP2α does not typically associate through CTCF/cohesin complexes but through less well defined sequences. It is therefore likely that multiple endogenous pathways may contribute to DNA break formation in HPV positive cells and that they play roles in regulating viral functions. Overall, our studies demonstrate critical roles for TOP2β in inducing DNA breaks and regulating the HPV life cycle.

## Materials and Methods

### Cell Culture and Antibodies

Normal human foreskin keratinocytes (HFKs) were isolated from de-identified neonatal specimens. HFK cells were transfected with HPV16 or HPV31 genomes to generate HFK16 and HFK31 cell lines that contain episomal HPV DNA as previously described (18). CIN612 cells were isolated from a cervical cancer biopsy and contain HPV31 episomes (19). All keratinocytes were co-cultured in E-Media with growth arrested NIH3T3 fibroblasts as described (3). Antibodies: phosphorylated CHK1 (pCHK1) (#12302; CST), phosphorylated CHK2 (pCHK2) (#2661; CST), FANCD2 (#100182; Novus), phosphorylated H2Ax (*γH2AX*) (#05636 Millipore), phosphorylated SMC1 (pSMC1) (#4805S; CST), BRCA1 (#OP92; Millipore), RAD51 (#NB100148; Novus),TOP2β (A300-950A; Bethyl Laboratories)

### Calcium Induced Differentiation

Keratinocytes were induced to differentiate through a calcium mediated switch (3). Briefly, keratinocytes were plated at 5 million cells per 10cm dish in M154 keratinocyte media (ThermoFisher) containing .07 mM Ca^2+^ and human keratinocyte growth serum (HKGS; ThermoFisher). The next day, the media was changed to .03mM Ca^2+^ in M154 containing HKGS. After 24 hours, the media was changed once again to M154 containing 1.5 mM Ca^2+^ in the absence of HKGS. These cells were then incubated in high calcium media for 72 hours to allow for proper differentiation.

### Western Blot Analysis

For western analysis of keratinocytes, cells were trypsinized from 10cm plates and the cell pellets were washed 3x in cold PBS. Cell pellets were then resuspended in an appropriate volume of RIPA buffer for lysis and incubated on ice for 15 minutes to facilitate lysis. Lysates were then quantified using a Pierce BCA protein assay kit to determine the protein concentration of each sample. Lysates were then combined with 6x SDS protein loading buffer and passed through a 25G syringe to homogenize lysates. Next, lysates were boiled at 100°C for 10 minutes to denature the samples. Samples were then centrifuged at 15,000 rpm for one minute and then were either directly frozen or used for Western blot analysis. For each western blot, 20μg of protein was loaded into each well of a pre-cast 10% polyacrylamide gel and run at 70V for 2 hours. Proteins were then transferred onto a PVDF membrane using wet transfer apparatus at 400 mA for 90 minutes at 4°C. The membrane was then blocked in 5% non-fat milk in TBST (Tris, NaCl, Tween) on a rotating platform at room temperature for 1 hour. Primary antibodies were then added to TBST and membranes were incubated overnight on a rotating platform at 4°C. The following day, membranes were washed 3x with TBST for 5 minutes each and were incubated with secondary antibodies conjugated with HRP for 1 hour at room temperature. After an additional 3x washes in TBST, membranes were activated using ECL blotting substrate and bands were visualized using a LICOR imaging station.

### Neutral COMET assay

Cells were seeded onto glass slides suspended in low melting agarose, lysed and electrophoresed according to manufacturer’s instructions (Trevigen Comet assay (ESII system). Comets were visualized using a Zeiss Axioscope and the percent tail DNA was quantitated using open source software OpenComet for FIJI.

### Southern Blot Analysis

Keratinocytes were first washed 3x in room temperature PBS and then trypsinized from 10cm plates and transferred to 15mL conical tubes. Cells were then pelleted and washed in PBS. Cell pellets were resuspended in 3mL Southern Lysis buffer. To each sample, RNAse A was added to a final concentration of 50μg/mL and incubated at room temperature for 15 minutes. Next, SDS was added to a final concentration of 0.2% along with Proteinase K to a final concentration of 50μg/mL and samples were incubated overnight at 37°C. The next day, samples were passed through an 18G syringe 5x and DNA was extracted using phenol/chloroform/isoamyl alcohol-based extraction. Following extraction, DNA was precipitated by adding 6mL of 100% ethanol to each sample and the samples incubated overnight at −20°C. The following day, precipitated DNA was pelleted by centrifugation at 5,000 rpm for 30 minutes at 4°C. DNA pellets were then washed 3x in 70% ethanol and were then air dried before resuspension in TE buffer. DNA was t quantified using a nanodrop UV spectrophotometer. For each sample, 5ug of DNA was digested using XhoI restriction enzyme. Digested DNA was separated on a 0.8% agarose gel at 40V for 16 hours. DNA was depurinated by washing the gel in HCl followed by neutralization in NaOH and transferred onto a nylon membrane using a vacuum apparatus. Transferred DNA was then neutralized in 2x sodium citrate buffer (SCC) and UV crosslinked to the nylon membrane. The membrane was then equilibrated in a pre-hybridization buffer before the addition of ^32^P labeled HPV31 DNA probe. Following addition of radiolabeled probe, the membrane was incubated overnight and washed in SCC the following day. After washing, DNA bands were visualized using autoradiography film.

### Exonuclease V assay

Episomal DNA was measured using an Exonuclease V assay described by the Sapp lab (27). Total DNA was isolated from cells using the phenol: chloroform followed by precipitation with ethanol. DNA was then quantitated using Qubit 4 fluorometer with the BR DNA assay kit (Invitrogen) and 1ug of DNA was digested using the exonuclease V. DNA was then purified using GeneJet PCR purification (Thermo Fisher) and amplified with Roche light cycler 480 using primers specific to E7. Forward primer 5’ATGAGCAATTACCCGACAGCTCAGA and reverse primer 5’ AGACTTACACTGACAACAAAAGGTAACGAT.

### q-RT-PCR for E7 and TOP2β transcripts

Total RNA was isolated from lysed cells using an EZ-10 spin column total RNA minipreps super kit (Bio Basic) following manufacture instructions. RNA was then digested with DNase1 and purified again through RNA mini prep kit again. cDNA was made using high capacity RNA-to-cDNA kit (Applied Biosystems) following manufacture instructions. cDNA was quantitated using Qubit 4 fluorometer with a ssDNA assay kit (Invitrogen). qRT-PCR reactions were performed using primers specific to E7 or TOP2b with a Roche light cycler 480. E7: forward primer 5’ATGAGCAATTACCCGACAGCTCAGA and reverse primer 5’ AGACTTACACTGACAACAAAAGGTAACGAT (IDT). Top2β: Forward primer 5’CAGCCCGAAAGACCTAAATAC and reverse primer 5’ATCTAACCCATCTGAAGGAAC obtained from Sigma-Aldrich.

### ChIP-qPCR

Keratinocytes were fixed in 1% formaldehyde in PBS for five minutes. Cells were then scraped from 10cm dishes and washed 3x in cold PBS. Fixed cells were then lysed by resuspending cells in 500μL of RIPA buffer on ice. Lysates were sonicated in 100 μL aliquots using a 30 second on/90 seconds off cycle for 20 minutes on high output. Sonicated lysates were then stored at −80°C. Protein G beads were added and pre-cleared overnight on a rotating platform using 0.5 mg/ml BSA and 0.5 mg/mL salmon sperm in PBS. The following morning, beads were magnetically separated and 2μg of antibody was added to each IP reaction followed by incubation for eight hours at 4°C on a rotator. Next, the beads were magnetically separated and 100 μL of sonicated cell lysate was added to each reaction along with 900μL of RIPA buffer. IP reactions were then incubated overnight at 4°C on a rotator. The following day, IP reactions were magnetically separated, and the beads were washed 8x in ice cold RIPA buffer. To elute captured DNA/protein, the beads were resuspended in an elution buffer (5mM EDTA, 1% SDS, 50 mM TRIS pH 7.5, 50 mM NaCl) and incubated for 30 minutes at 55°C with shaking. The beads were then separated by centrifugation at 15,000g and the supernatant was moved to a clean Eppendorf tube. The eluted fractions were next reverse crosslinked at 55°C overnight. Immunoprecipitated DNA was then purified using standard QIAGEN quick purification columns. DNA was quantified by qPCR using primers specific to HPV genomes as well as Alu repeat sequences specific to human genomic DNA. The relative amount of specific DNA per sample was normalized to an input chromatin fraction that did not undergo immunoprecipitation.

### Immunofluorescence, microscopy and Image J analysis

Cells were grown on treated coverslips under media. Prior to analysis, the cells were fixed with 4% paraformaldehyde, permeabilized in PBS with Triton-X100 (PBST), blocked with Normal Goat Serum (NGS), and probed with primary antibodies in NGS at 4°C overnight. After three washes in PBST, slides were incubated with secondary antibodies followed by DAPI incubation, and coverslips mounted. Images were taken on a Zeiss Axioscope and the data were imported into ImageJ for analysis. The percentage of positive-staining cells was evaluated by comparing the number of cells with nuclear focal staining to the total number of cells as determined by DAPI staining and greater than 20 cells were examined in each group.

### Statistical analysis

IBM SPSS Statics24 software was used for statistical analysis. Comparison between two groups was performed with unpaired Student’s t test. P values of <0.05 were considered to be statistically significant.

## Acknowledgements

LAL was supported by grants from the National Cancer Institute (RO1CA059655 and RO1CA142861). We thank Dr. Elona Gusho for help with analysis and figures.

